# Evolutionarily-conserved behavioral plasticity enables context-dependent performance of mating behavior in *C. elegans*

**DOI:** 10.1101/2023.04.26.538441

**Authors:** Vladislav Susoy, Aravinthan D.T. Samuel

**Author notes:** Correspondence (VS); (AS).

## Abstract

Behavioral plasticity helps humans and animals to achieve their goals by adapting their behaviors to different environments. Although behavioral plasticity is ubiquitous, many innate species-specific behaviors, such as mating, are often assumed to be stereotyped and unaffected by plasticity or learning, especially in invertebrates. Here, we describe a novel case of behavioral plasticity in the nematode *C. elegans* – under a different set of naturalistic conditions the male uses a unique, previously undescribed set of behavioral steps for mating. Under standard lab conditions (agar plates with bacterial food), the male performs parallel mating, a largely two-dimensional behavioral strategy where his body and tail remain flat on the surface and slide alongside the partner ‘s body from initial contact to copulation. But when placed in liquid medium, the male performs spiral mating, a distinctly three-dimensional behavioral strategy where he winds around the partner ’s body in a helical embrace. The performance of spiral mating does not require a long-term change in growing conditions but it does improve with experience. This experience-dependent improvement involves a critical period – a time window around the L4 to early adult stage, which coincides with the development of most male-specific neurons. We tested several wild isolates of *C. elegans* and other *Caenorhabditis* species and found that most were capable of parallel mating on surfaces and spiral mating in liquids. We suggest that two- and three-dimensional mating strategies in *Caenorhabditis* are plastic, conditionally expressed phenotypes conserved across the genus, and which can be genetically “fixed ” in some species.

## Results

We sought to investigate the mating performance of male *C. elegans* in different naturalistic conditions. When kept in standard lab conditions on agar plates with bacterial food, the male performs parallel mating, where his movements are confined to the two-dimensional agar surface and his tail remains parallel with his partner ‘s body from contact to copulation (Figure 1A). We discovered that when *C. elegans* males are placed n liquid hydrogel (Muschiol and Traunspurger, 2007; Gilarte et al., 2015), the male does not perform parallel mating. Instead, upon contact with the hermaphrodite, the male wraps his body around her (Figure 1B). This behavior resembles spiral mating, which has been observed in other nematode species. Obligate spiral mating has been previously reported in only four *Caenorhabditis* species (Kiontke et al., 2011; Stevens et al., 2019), and both parallel and spiral mating have been observed in *C. remanei* (Sudhaus and Fitch, 2001). It has been suggested that spiral mating evolved twice in *Caenorhabditis* (Stevens et al., 2019). Our results show that parallel and spiral mating strategies rep-resent behaviorally plastic phenotypes that can be induced in *C. elegans* and other species by a simple environmental change.

**Figure 1.**
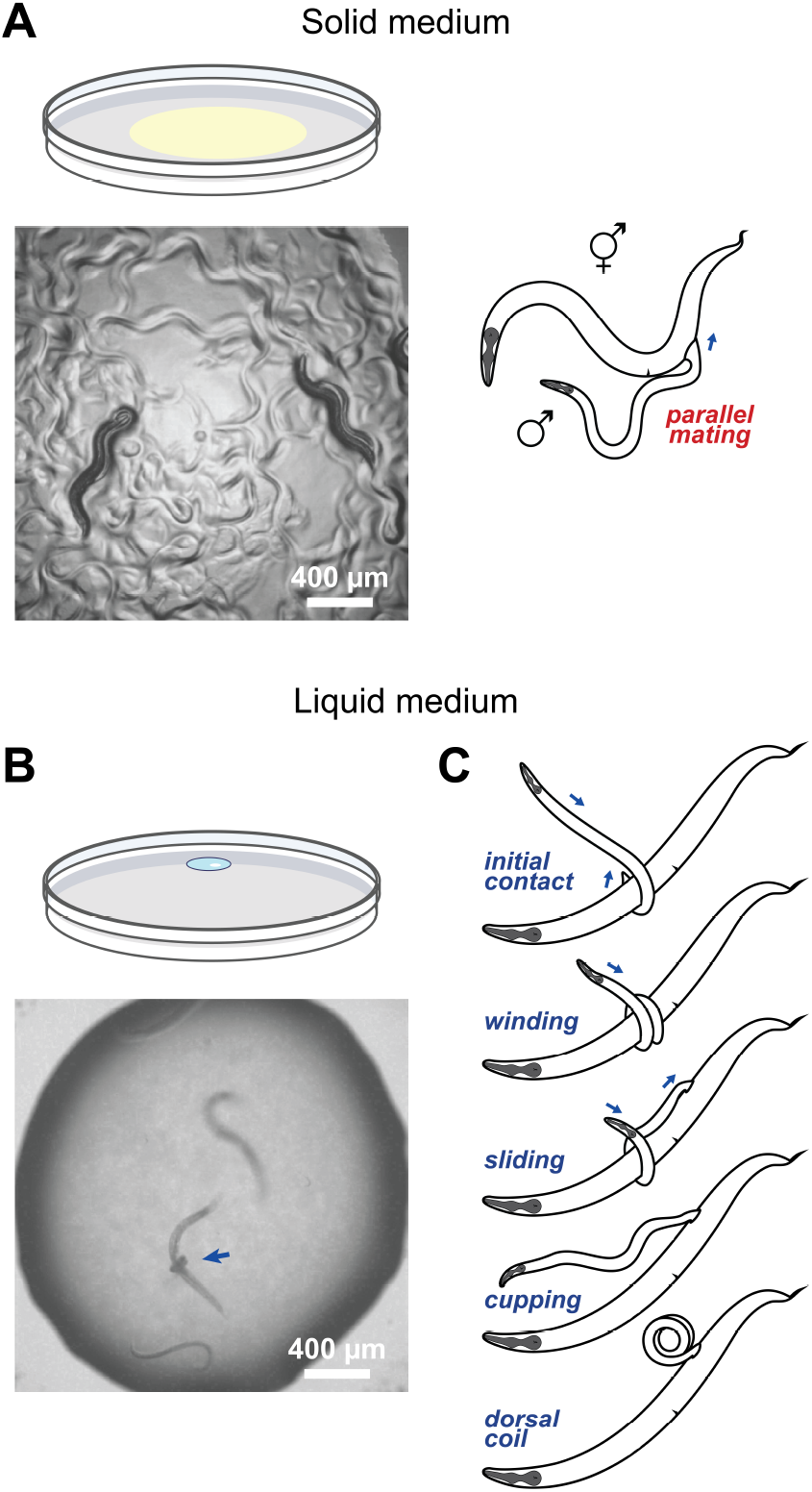
Behavioral plasticity in *C. elegans*. *C. elegans* males use two different strategies for mating. On solid surfaces the male performs parallel mating, **(A)** whereas in liquid the male uses spiral mating **(B)**. Blue arrow points at a male *C. elegans* wrapped around a hermaphrodite. **(C)** Spiral mating in liquid usually involves a stereotyped sequence of behavioral steps or motifs.

Video recordings of spiral mating of multiple *C. elegans* males revealed that this behavior involves a stereotyped sequence of behavioral steps or motifs (Figure 1C, Video S1). First, initial contact of the male tail with the hermaphrodite induces a sharp ventral flexion of the tail. When this first maneuver is successful, the male grips the hermaphrodite with his tail. In the second step, the male winds himself around the hermaphrodite, forming a helical coil around her. In the third step, the male maintains his helical hold of the hermaphrodite with his anterior end, which allows him to slide his tail backwards to a new location on the hermaphrodite. As the male tail slides backwards, he unwinds his anterior end. When he is completely unwound, the male only holds the hermaphrodite by his tail, which cups onto her body. Occasionally, in the fourth step, the male bends his entire body dorsally, forming a dorsal coil. The last three steps of spiral mating can be repeated multiple times in sequence, until the male successfully cups onto the vulva with his tail and completes copulation or until he loses the hermaphrodite. The performance of spiral mating resembles the concertina locomotion used by arboreal snakes to climb trees (Byrnes and Jayne, 2014). In the male *C. elegans*, spiral mating might help to mate in liquid environments where the low substrate resistance and the three-dimensional nature of the task pose challenges for parallel mating. We conclude that spiral mating is a complex behavior consisting of separate behavioral motifs; it is distinct from parallel mating observed on solid surfaces, and likely represents an adaptation to liquid environments.

Next we tested whether males can successfully mate in liquid and produce progeny. We used males and hermaphrodites that carried a mutation in *fog-2*, which prevented hermaphrodites from self-fertilizing. We divided males into two groups: one group raised with hermaphrodites on agar plates, the other group raised with hermaphrodites in liquid. We tested the ability of both groups of males to mate with virgin hermaphrodites in liquid by counting progeny. The males that were raised in liquid from egg to adult stages were able to mate successfully in liquid (Figure 2A). In contrast, the males that were raised on agar plates and only transferred to liquid to mate as adults failed to produce any progeny. We conclude that (i) *C. elegans* males can mate successfully in liquid, and (ii) this ability depends on prior experience.

**Figure 2.**
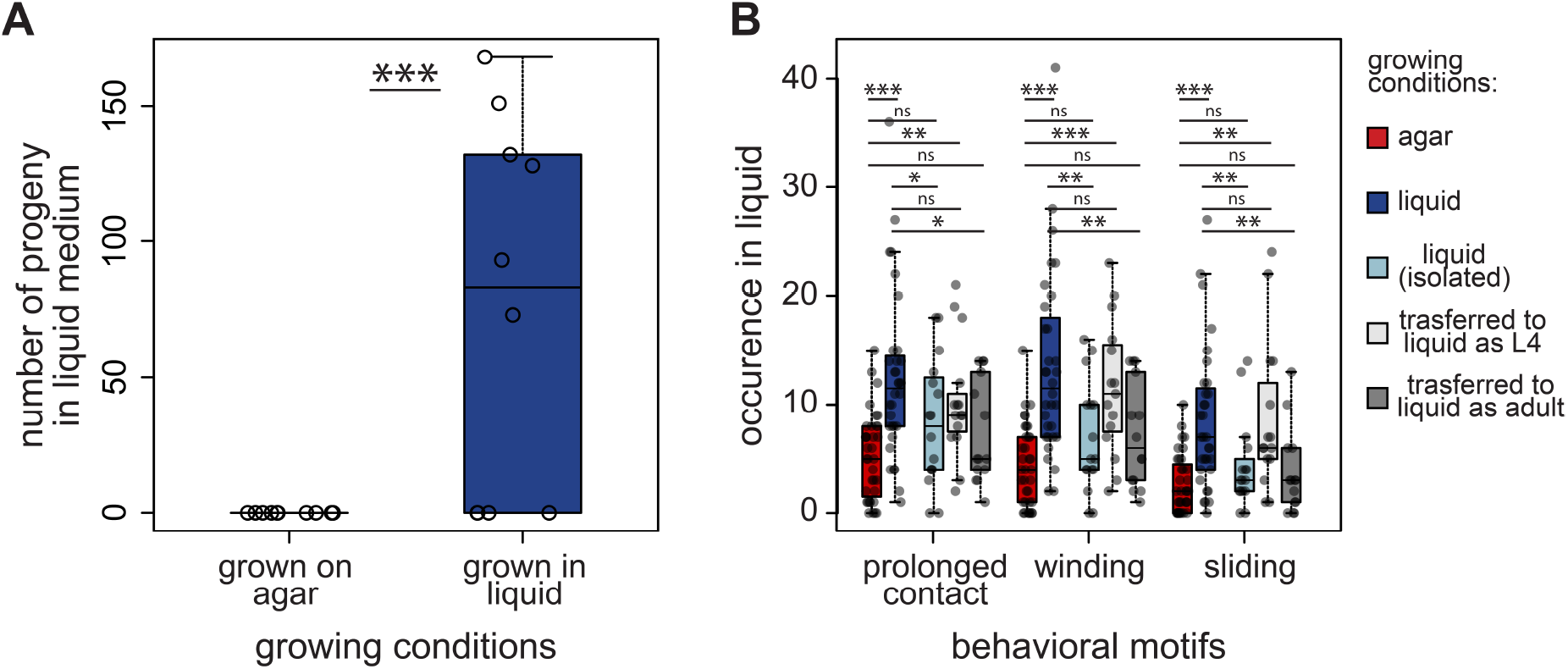
The performance of spiral mating improves with experience. **(A)** Males can mate successfully in liquid medium and produce progeny but only if they have been grown in liquid from the egg stage. The number of progeny per mating test is shown; mating tests were performed in liquid. **(B)** The performance of individual steps of spiral mating improves with experience. The following motifs are shown: (i) prolonged contact – tail contact with the hermaphrodite lasting >10 sec; (ii) winding – winding of the male around the hermaphrodite (iii) sliding - sliding of the male tail backwards along the hermaphrodite body. t-test, * *P* < 0.05, ** *P* < 0.01, *** *P* < 0.001. Boxplots show the median, Q1, and Q3 values; whiskers extend to a maximum of 1.5 IQR beyond the box.

As mating success is influenced by prior experience, we wanted to know if the performance of individual steps of spiral mating differed between the males grown in liquid and on agar plates. The males raised on agar plates and transferred to liquid as adults were able to attempt all individual steps or spiral mating. However, the males raised in liquid performed all steps of spiral mating with significantly greater frequency, displaying more instances of prolonged contact with the hermaphrodite, winding around her, and sliding of the tail (Figure 2B). The ability of male *C. elegans* to initiate spiral mating does not require long-term experience in liquid environments, but the execution of spiral mating improves with time spent in liquid medium, which is indicative of learning.

We asked whether experience with the liquid environment is sufficient for improving the performance of spiral mating behavior or whether direct experience with hermaphrodites in the liquid environments might also matter. We raised individual males in isolation in separate droplets. When males raised in isolation were introduced to hermaphrodites in liquid environments, they exhibited fewer instances of all three motifs of spiral mating (prolonged contact, winding, and sliding) compared to the males raised in liquid environments in the presence of hermaphrodites (Figure 2B). This suggests that males improve their spiral mating behavior with actual mating experience, not just by exposure to a liquid environment.

Many instances of long-term behavioral plasticity that are induced by life experience involve critical periods, developmental intervals when plasticity and learning are potentiated (Wiesel and Hubel, 1963; Lorenz, 1937; Doupe and Kuhl, 1999). To investigate whether experience-dependent improvement in mating performance involves a critical period, we transferred males between agar plates and liquid medium at different points in development. In particular, we focused on the time around the L4 larval stage when many neurons of the male-specific mating circuit are born (Sulston et al., 1980). We found that males transferred to liquid as L4 larvae showed similar performance of spiral mating compared to the males cultured in liquid from the egg stage. In contrast, males grown on agar and transferred to liquid overnight as adults showed similar performance of spiral mating to the males cultured on agar (Figure 2B). These results indicate that behavioral plasticity is potentiated around L4-early adult stages.

We next tested whether this new case of behavioral plasticity is conserved across different wild isolates of *C. elegans*. We se-lected six genetically distinct isolates and tested if they can perform spiral mating when placed in liquid medium (Figure 3A). When cultured in liquid from the egg stage, all isolates except QX1211 were able to perform spiral mating. We also tested whether the ability to perform spiral mating in liquid medium is conserved across different *Caenorhabditis* species. We have selected eight species from different groups of *Caenorhabditis*: *C. sulstoni, C. inopinata, C. remanei, C. sinica, C. brenneri, C. utelia, C. virilis*, and *C. monodelphis*. All tested species were able to perform both parallel mating on solid surfaces and spiral mating in liquid, in a manner similar to *C. elegans* (Figures 3B– 3C). We conclude that spiral mating is a plastic behavioral phenotype conserved across genetically distinct wild isolates of *C. elegans* as well as across species of *Caenorhabditis*.

**Figure 3.**
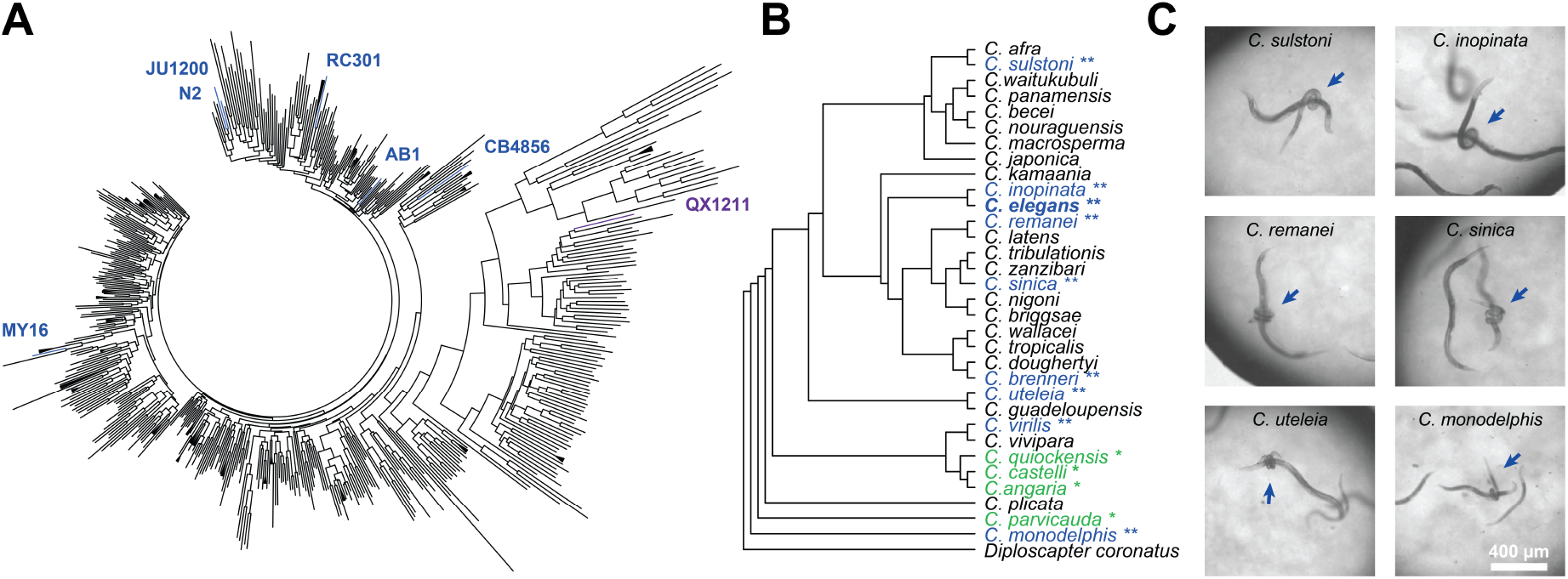
Spiral mating is conserved across isolates and species. **(A)** Six genetically distinct wild isolates of *C. elegans* were selected for mating tests. All isolates except QX1211 were able to perform spiral mating when placed in liquid. Phylogeny of the *C. elegans* wild isolates is based on Cook et al., 2017 (Cook et al., 2017) **(B)** All species of *Caenorhabditis* are known to perform parallel mating on agar, except four species in which spiral mating is obligate (indicated with *). All nine species that were tested for spiral mating in liquid (indicated with **) were able to perform it in a manner similar to *C. elegans*. The remaining species have not been tested for spiral mating. Phylogeny of *Caenorhabditis* species, adapted from Stevens et al., 2019 (Stevens et al., 2019). **(C)** Still frames from video recordings showing spiral mating of six *Caenorhabditis* species. Blue arrows point at the males wrapped around their partners.

Plastic behavioral phenotypes are often accompanied by correlated morphological changes (Pfennig, 1990; Moczek and Emlen, 2000; Vijendravarma et al., 2013; Wilecki et al., 2015). We used geometric morphometrics to compare the morphologies of male tail sensilla between males grown on agar plates and in liquid medium (Figure 4A). Principal component analysis (PCA) of the tail shape and form (shape+size) showed that males raised on solid vs. liquid media tend to occupy different regions of the Procrustes morphospace (Figure 4B). Detailed analysis showed that four pairs of male rays – 2, 3, 4, and 6 – are especially affected (Figure 4C). Rays 2, 4, and 6 are shorter and thicker in males grown in liquid whereas ray 3 is also more curved in the middle. Previous studies revealed that these rays play key roles in hermaphrodite recognition and scanning Liu and Sternberg (1995); Barr and Garcia (2006); Jarrell et al. (2012); Susoy et al. (2021). Although the functional significance of the observed context-dependent changes in ray morphology remains to be characterized, we hypothesize that these changes might facilitate mating in different conditions.

**Figure 4.**
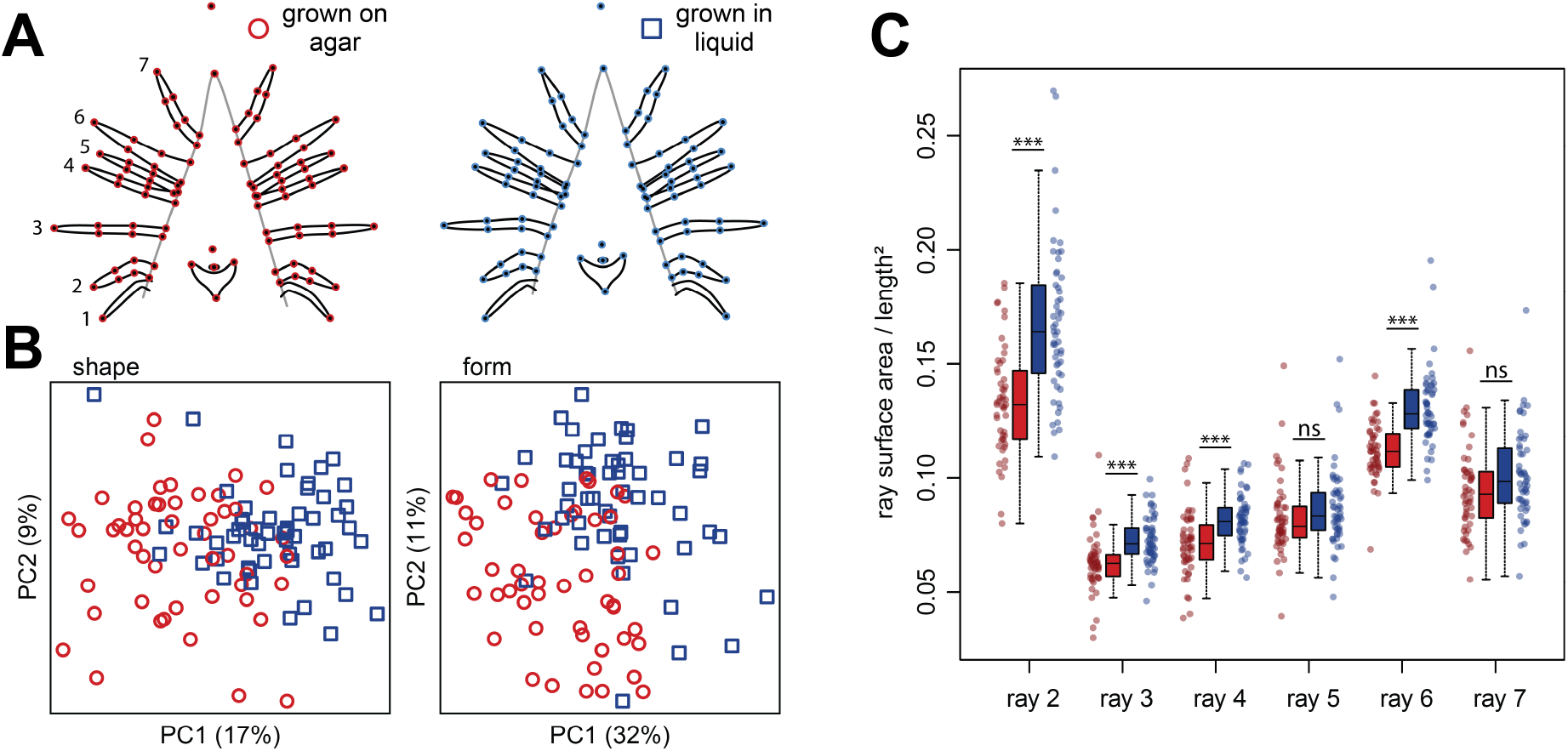
Different sensilla morphologies in different media. When cultured in liquid medium, males show a morphological change in the shape and form of their tail sensilla. **(A)** Average shapes of tail and sensilla structures in males grown on agar (left) and in liquid (right). Individual dots show morphological landmarks (open circles) and semi-landmarks (closed circles) with the black lines demarcating male tail rays 1-7 and tail structures. **(B)** Males grown on agar (red circles) and in liquid medium (blue squares) tend to occupy different subspaces of the Procrustes morphospace of shape (left) and form (shape+size, right). **(C)** Males grown in liquid (blue) have shorter and thicker sensory rays than males grown on agar (red). t-test, * *P* < 0.05, ** *P* < 0.01, *** *P* < 0.001. Boxplots show the median, Q1, and Q3 values; whiskers extend to a maximum of 1.5 IQR beyond the box.

## Discussion

Our results show that a simple environmental shift induces a dramatic change in what has been considered to be a stereotyped species-specific behavior in *C. elegans* – mating. When cultured on solid surfaces and in liquid, *C. elegans* males adopt different behavioral strategies – parallel and spiral mating – each with its own set of unique behavioral motifs.

This conditional change in mating behavior involves both contextual plasticity, where the animal shows an immediate behavioral response to a new environment, as well as longterm developmental plasticity, where behavioral performance improves with experience. The fact that naïve males grown on solid surfaces are able to perform spiral mating shortly after being placed in liquid suggests that the existing neuronal circuit can accommodate both behaviors. Nevertheless, the performance of spiral mating improves significantly when the male is raised in liquid in the presence of hermaphrodites from an early stage. The developmental mechanisms behind this long-term plasticity may be complex and involve changes in neuronal circuit wiring, gene expression, muscle system development, and morphology.

The ability to shift between parallel and spiral mating is conserved across different wild isolates of *C. elegans* and different *Caenorhabditis* species. Such conservation of plastic traits is not uncommon in closely related species with similar ecological niches, as they are likely to encounter similar environmental fluctuations. For example, many species of diplogastrid nematodes develop either narrow-mouthed bacterivorous morphs when bacterial food is abundant or wide-mouthed predatory morphs when bacterial food is scarce (Susoy et al., 2015). In some taxonomic groups, formerly environmentally-induced plastic traits can become “fixed “ via genetic assimilation (i.e., expressed unconditionally in all environments) (Waddington, 1953; West-Eberhard, 2003). In the genus *Caenorhabditis*, four species exclusively exhibit spiral mating, even when cultured on solid media. This behavioral specialization is also accompanied by morphological changes in the male sensilla (Kiontke et al., 2011). Given the prevalence of the context-dependent plasticity in mating behavior across the genus – including in *C. monodelphis*, the outgroup for all other *Caenorhabditis* spp. – we speculate that these four species might have lost their ancestral plasticity, which accords with the idea that behavioral plasticity and learned traits can precede the evolution of genetically fixed traits (West-Eberhard, 2003).

## Methods

### Resource Availability

#### Lead contact

Further information and requests for resources and reagents should be directed to Aravinthan D.T. Samuel

#### *Caenorhabditis* strains

CB4088 *him-5 (e1490) V*

CB4108 *fog-2 (q71) V*

AB1 *C. elegans* wild isolate

CB4856 *C. elegans* wild isolate

JU1200 *C. elegans* wild isolate

MY16 *C. elegans* wild isolate

RC301 *C. elegans* wild isolate

QX1211 *C. elegans* wild isolate

JU2788 *C. sulstoni* wild isolate

NKZ35 *C. inopinata* wild isolate

JU1082 *C. remanei* wild isolate

JU1201 *C. sinica* wild isolate

JU2585 *C. utelia* wild isolate

PB2801 *C. brenneri* wild isolate

JU1968 *C. virilis* wild isolate

JU1667 *C. monodelphis* wild isolate

### Culture media

For the standard cultures, we used 6 cm NGM agar plates seeded with *E. coli* OP50 bacteria. For the hanging drop method, we used 0.0015% gellan gum in the NGM buffer, adapting protocols by Muschiol and Traunspurger (2007) and Gilarte et al. (2015). Before use, approximately one half of an *E. coli* OP50 lawn was scraped from a single seeded NGM plate and mixed well with 500 μL of the gellan gum medium. The resulting suspension had a semi-liquid consistency, where the nematodes could swim freely without sinking. We refer to this medium as “liquid medium “. Drops of the liquid medium were applied to the underside of the lids of unseeded NGM petri dishes, and the nematodes or their eggs were transferred to the hanging drops.

### Nematode conditioning

To test how growing conditions affect mating performance, male nematodes were conditioned using the following regimes: (i) 20 eggs were placed on an agar plate on day 1, and the adult males were used for behavioral experiments on day 5; (ii) 20 eggs were placed in 50 μL of liquid medium on day 1, the cultures were checked on day 4, and if the food was low, the nematodes were picked into fresh medium; the adult males were used for behavioral experiments on day 5; (iii) the nematodes were cultured on agar from the egg stage and transferred to 50 μL liquid medium on day 3 at the L4 stage; (iv) the nematodes were cultured on agar from the egg stage, and were transferred to liquid medium overnight on day 4 at the adult stage; (v) to grow males individually in isolation from other males and hermaphrodites, single eggs were placed inside 10 μL hanging drops. All culture plates were sealed with Parafilm M. Behavioral recordings were performed on day 5. To test for spiral mating among *C. elegans* wild isolates and *Caenorhabditis* species, nematodes were cultured in liquid from the egg stage, except *C. monodelphis*, which were transferred to liquid as L4 larvae. Males were always paired with females/ hermaphrodites of the same strain/ species.

### Mating tests

Two adult *fog-2* males (day 4) and two adult *fog-2* hermaphrodites were placed in 10 μ L of liquid medium, and allowed to mate overnight. The males were removed from the medium, while the hermaphrodites were allowed to lay eggs. The number of progeny was counted four days later.

### Mating tests – video recordings

Two males were placed together with three hermaphrodites inside a 4 μL hanging drop. The males were picked from either solid or liquid medium, depending on the conditioning regime (see above). The hermaphrodites were always picked from agar plates, to minimize any condition-dependent variability in the hermaphrodites ‘ behavior. The behavior of the males and hermaphrodites was recorded using a Grasshopper3 camera at 5 fps for 30 minutes. The recordings were analyzed blindly. We tabulated the occurrence of the following behavioral motifs: (i) prolonged contact of the male tail with the hermaphrodite (>4 seconds); (ii) wrapping of the male around the hermaphrodite (occasionally, the male formed a loose coil, using his anterior end as a lever to push his tail backwards) (iii) sliding of the tail backwards, which followed the wrapping around the hermaphrodite.

### Morphology

To test whether males cultured on agar and in liquid had morphological differences in their tail structures, we used geometric morphometrics, adapting an approach previously used for the nematode stomatal structures (Susoy et al., 2015). Briefly, nematodes were picked from their culture medium and mounted on a glass slide using a 10% agar pad and a 2–3 μL drop of the liquid medium such that the ventral side of the male tail was pressed against the cover slip. Images of the tail were taken with a Nikon microscope in DIC mode, using a 100 × 1.45 NA oil objective, CoolSnap camera, and μManager software. Males showing gross morphological abnormalities (missing rays, etc.) were excluded from further analyses. We assigned 45 landmarks and 48 semilandmarks to homologous tail structures using Fiji (Schindelin et al., 2012), as illustrated in Figure 4A. Ray 1 was only assigned a single landmark at its distal tip due to the uncertainty in placing landmarks at its base. Rays 8 and 9 were excluded completely because they often fused into one structure. The landmarks were used for Procrustes analysis with the R package “geomorph ” (Adams et al., 2013). We then performed PCA on the Procrustes-superimposed landmark coordinates. For the PCA of shape, we used landmark positions only. For the PCA of form, we additionally included the logarithm-transformed centroid size, accounting for size as well as shape differences. Additionally, for each ray, we quantified the ray surface area/midline length^2^ ratio using the landmark coordinates. A larger ratio indicates shorter and thicker rays.

## Supplemental Information

**Video S1. Spiral mating in male *C. elegans***. When placed in liquid medium, *C. elegans* males perform spiral mating, a three-dimensional behavioral strategy where he winds around the hermaphrodite. Spiral mating is different from parallel mating, a largely two-dimensional strategy typically observed under standard lab conditions – agar plates with bacterial food.

## Acknowledgements

We thank David Zimmerman for his comments on the manuscript. We thank CGC, which is funded by NIH Office of Research Infrastructure Programs (P40 OD010440) for providing strains. This work was supported by NIH (R01 NS113119).

## Author Contributions

VS conceived the project, performed the experiments, and analyzed the data. VS and AS wrote the manuscript.

